# DNA-polymerase guided elimination of paternal mitochondrial genomes: An escape-proof obstacle to their transmission

**DOI:** 10.1101/086702

**Authors:** Zhongsheng Yu, Patrick H. O’Farrell, Nikita Yakubovich, Steven Z. DeLuca

**Affiliations:** Department of Biochemistry and Biophysics, UCSF, San Francisco, CA; Current address: Carnegie Institution of Science, Dept. of Embryology, Baltimore, MD

## Abstract

Mitochondrial DNA is predominantly inherited from only one parent. In animals this is usually the mother. This program is not in the interest of the paternal mitochondrial genome whose potential to contribute to future generations is restricted. However, in a dramatic example of genetic conflict, nuclear programs ensure the outcome. Two large mitochondria extend the length of *Drosophila* sperm tails. The hundreds of nucleoids in these mitochondria vanish during spermatogenesis eliminating their potential for transmission. Our previous work showed that mutational inactivation of EndoG, a nuclear encoded mitochondrial endonuclease, slows elimination of mitochondrial genomes. Here, we show that knockdown of the nuclearly encoded mitochondrial DNA polymerase, Tamas, produces a much more complete block of mtDNA loss. Recruitment of Tamas to the nucleoid at the time of its disappearance suggests a direct contribution to the elimination, but the 3′-exonuclease function of the polymerase is not needed. While DNA elimination is a surprising function for DNA polymerase, its use to restrict paternal genomes provides a strategy that cannot easily be evaded by the mitochondrial genome without compromising its replication.

## Introduction

In a wide range of species, mitochondrial DNA is only transmitted from one parent [1,2]. This phenomenon, familiar to many people as maternal inheritance or as cytoplasmic inheritance, is often thought of as a passive consequence of the disproportionately large cytoplasmic volume of the female gamete. However, uniparental inheritance extends to organisms in which both gametes are a similar size, indicating a need for other explanations [3,4]. Two categories of active mechanisms have been described. In one the DNA is eliminated within the mitochondria from one of the two sexes [5–7], and in the other, the mitochondria themselves, and presumably any residual DNA, are eliminated [8–10]. In *Drosophila*, both the DNA and the mitochondria are targeted. The mitochondrial DNA is eliminated in the male during spermatogenesis, as part of developmental programs that restructure the mitochondria during the formation of the sperm tail (Figure 1) [7]. Though already devoid of genomes, the large sperm mitochondria that enter the zygote upon fertilization are subsequently destroyed [11,12]. It appears that multiple tiers of regulation create a formidable barrier blocking transmission of mitochondrial DNA from one of the two sexes, usually the male.

The existence of active mechanisms enforcing uniparental inheritance suggests that it is biologically important to avoid biparental inheritance. Theory and experiment argue that biparental inheritance would allow mitochondrial genomes to pursue an infectious lifestyle: because biparentally inherited mitochondrial genomes that gain a transmission advantage (e.g. a mutant having enhanced replication) would be passed on to all progeny from either parent, they would have a transmission advantage and could spread infectiously through an entire interbreeding population [13–17]. Such an infectious spread of mitochondrial genomes favors selfish mutations, which can succeed even when associated with mutations detrimental to the host [15]. In contrast, uniparental/maternal inheritance limits juggernaut genomes to clonal female (matroclinous) lineages that will fail unless function is maintained. In this way, uniparental inheritance aligns the interests of extranuclear genomes with the fate of the organism and substantially suppresses conflicting evolutionary pressures that would otherwise disrupt the collaboration between nuclear and mitochondrial genomes [18].

But uniparental inheritance creates another conflict. If the host develops an active program to prevent transmission of paternal mitochondrial DNA, mitochondrial DNAs are likely to acquire mutations that evade the restriction imposed on it. Indeed, biology provides examples where the restriction is bypassed. Bypass is particularly clear when unipartental inheritance of the mitochondrial genome is operative, but a second mitochondrial DNA element, a mitochondrial plasmid, achieves biparental transmission and spreads preferentially [19,20]. For example, the mF plasmid DNA of *Physarum polycephalum* evades mitochondrial DNA destruction and manipulates mitochondrial fusion to ensure its biparental inheritance thereby promoting its spread [20,21]. In another biological variant, it is the paternal mitochondrial genome that escapes the restriction to achieve transmission. A number of bivalve molluscs exhibit “doubly uniparental inheritance” in which male genomes are transmitted. But transmission is only to male progeny, and within the male progeny these paternal mitochondrial genomes only contribute to the germline: this effectively maintains a separation between the male and female mitochondrial lineages, but without the block to paternal mtDNA transmission [22–24]. With selection for transmission of paternal genomes continuously challenging the stability of maternal inheritance, the programs that restrict paternal genome transmission will be those that are not easily evaded.

Our analysis of the genetic basis for elimination of mitochondrial genomes during male gametogenesis in *Drosophila* suggests a strategy that would be difficult to evade by changes in the mitochondrial genome. We have found that the mitochondrial DNA polymerase is required for this DNA elimination program. If the host dictates whether the polymerase participates in a replication machine or a destruction complex, which both interact with the mitochondrial genome in the same way, the mitochondrial genome cannot evade the destruction complex without losing interaction with the replication machinery. This strategy could contribute to the enslavement of mitochondrial genome for the host’s purposes. In addition to enforcing maternal inheritance, mitochondrial genome elimination might occur at other stages, and its dysregulation could account for the finding that losses and disruptions of mitochondrial DNA occur in postmitotic tissues in several diseases that have been attributed to mutations in the mitochondrial DNA polymerase and its accessory proteins [25–27].

## RESULTS

### Mitochondrial DNA is eliminated during sperm development

*Drosophila* sperm originate from germ-line stem cells (GSCs) at the apical end of testes. Asymmetric divisions of GSCs generate another stem cell and a gonialblast that initiates spermatogenesis (Figure 1). The gonialblast undergoes four mitotic divisions with incomplete cytokinesis to produce 16 interconnected pre-meiotic cells, which, after a period of growth and preparation, undergo meiosis to produce a cyst of 64 interconnected spermatids. Just prior to the cellular reconfiguration that shapes the sperm, the numerous mitochondria fuse forming a large ball, the nebenkern, next to each nucleus of the cyst. The coiled appearance of this ball gives rise to the name onion-stage. This ball of mitochondria subsequently unfolds into precisely two giant mitochondria that bracket each centrosome and elongate dramatically in association with the axonemes to extend the sperm tails, which will reach the seemingly excessive length of 1.8 mm [28,29]. After completion of elongation, a specialized actin based structure called the individualization complex forms near spermatid nuclei (Figure 1). This structure then invests the axoneme and physically travels apically down the axoneme sweeping out excess cytoplasm, trimming the mitochondria, and remodeling the membranes to generate 64 individual sperm from the cyst [28]. In our previous work, we imaged foci of DAPI staining representing the compact DNA of the nucleoids within the mitochondria of developing spermatids, and found that nucleoids disappeared from the persisting mitochondria as the sperm tails approached their full length ([7] and Figure 2C). This destruction of nucleoids was promoted by a mitochondrial endonuclease, EndoG. Nonetheless, nucleoid elimination still occurred in mutants of this gene, though more slowly so that it was completed later during individualization — a stage marked by the moving individualization complexes that create a cystic bulge that travels the length of the bundle of tails (Figure 2D-E and see DeLuca and O’Farrell, 2012 for details).

**Figure 1.**
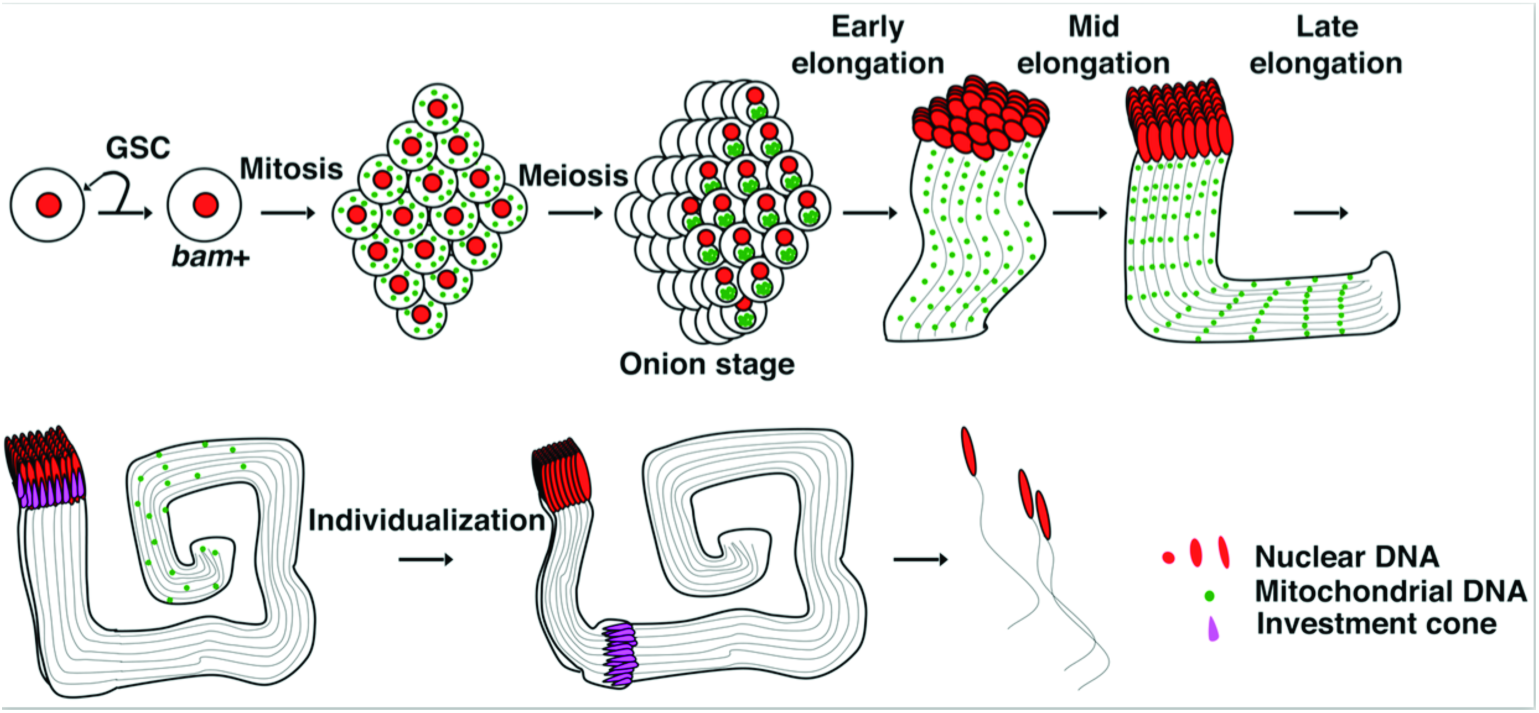
Coordination of mtDNA elimination with *Drosophila* spermatogenesis. Schematic of spermatogenesis with nuclei in red, mitochondrial genomes in green and the investment complex in magenta. The incomplete mitotic and meiotic divisions create a cyst of 64 spermatids that develop coordinately extending their tails as a bundle with 64 axonemes and 128 elongating mitochondria whose genomes disappear in proximal to distal wave as the tails reach full length prior to the individualization stage. We use tail length to define early, mid and late stages of elongation (10-1200µm, 1200-1700µm and >1700 µm, respectively).

### Tam is required for paternal mtDNA elimination

We modified and expanded our original candidate genetic screen in the hopes of defining mutations that more completely block the disappearance of mitochondrial genomes during spermatogenesis. We considered any gene having a known or predicted nuclease domain since the disappearance of DNA staining in the mitochondria would seem to require such an activity. In the current study, we used a driver of gene expression specific for early gametogenesis, *bam-GAL4*, to express *UAS-RNAi* to knockdown (KD) candidate genes in differentiating spermatids (Figure 1, *bam*+). This germline specific KD allowed us to assay phenotypes in spermatogenesis even if the candidate genes were needed earlier in development.

**Figure 2.**
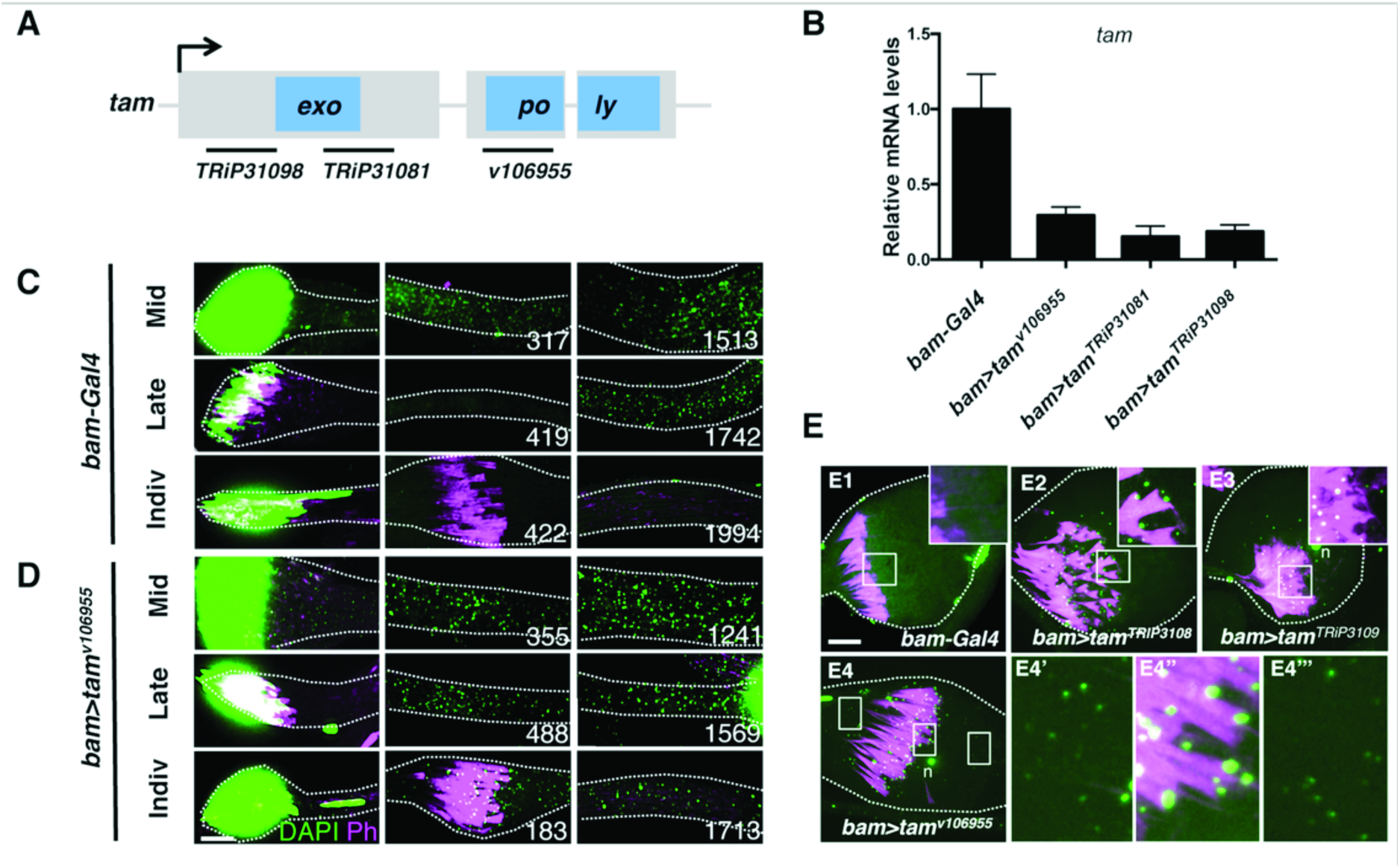
Knockdown of the mitochondrial DNA polymerase prevents nucleoids loss in developing spermatids. **(A)** Schematic of the *tam* gene showing regions encoding the exonuclease and polymerase domains and regions targeted by 3 non-overlapping RNAi hairpins used in this study. **(B)** qPCR measuring *tam* mRNA in *bam>tam^RNAi^* adult testis relative to control. Error bars represent the standard deviation for three independent experiments. **(C-E)** Spermatid bundles stained for DNA with DAPI (green) and actin with Phalloidin (Ph: magenta). Numbers indicate position of the image along the bundle of tails in microns. **(C)** *bam-GAL4* control. The full length of the cysts at the successive stages was 1590µm, 1864µm, and 2083µm. (D) *bam>tam^v106955^*. The full length of the cysts at successive stages was 1471µm, 1784µm, and 1803µm. **(E)** Cystic bulges showing that nucleoids are eliminated in the control (*bam-GAL4*) but not when Tam was knocked down by each of three RNAi constructs. The area within the rectangles with white borders are shown enlarged. E4 shows that nucleoids persist behind the investment cones, which move distally (to the right). n indicates the nuclear DNA fragment generated during individualization process. Scale bars are 10µm. Mid-, late-elongation and individualization stages are indicated as Mid, Late and Indiv.

The mitochondrial DNA polymerase, Pol gamma alpha, is encoded by the nuclear gene *tamas* (*tam*) [30,31]. We examined the possible role of the Tam protein in mtDNA elimination, because, in addition to its C-terminal polymerase domain (poly), it includes an N-terminal exonuclease domain (exo), which is thought to be important for replication-coupled proofreading (Figure 2A) [32–34]. Additionally, it had been observed that overproduction of Tam resulted in an mtDNA depletion phenotype [35]. When we expressed any of three non-overlapping transgenic RNAi hairpins targeting *tam* mRNA under *bam-GAL4* control (*bam*>*tam^RNAi^*), we observed persisting nucleoids in individualizing spermatids (Figure 2D and E). We verified that all three RNAi constructs expressed in differentiating germline cells of the testis greatly reduced tam mRNA in whole testis (Figure 2B). Because nucleoids are normally eliminated before spermatid individualization begins (e.g. Figure 2C), the abundant presence of nucleoids in individualizing *bam*>*tam^RNAi^*spermatids (Figure 2D and 3C) shows that Tam KD blocks normal nucleoid elimination.

### Tam plays more robust roles than EndoG during mtDNA elimination

The passage of the individualization complex trims the mitochondria and vesicles of mitochondrial material accumulate in the cystic bulge [36,37]. Previously, we found the cystic bulge also collected the low level of persisting mitochondrial nucleoids in the *EndoG* mutant, eventually eliminating them in the distal waste bag [7]. This appears to be a second mechanism that backs up the earlier elimination of mtDNA by DNA destruction (Figure 3C, *EndoG*). In spermatids from *bam*>*tam^RNAi^* flies, we also observed a higher density of nucleoids within the cystic bulge than in regions immediately in front of, or behind it, indicating that the cystic bulge also retains some capacity to collect nucleoids (Figure 2E). However, many mitochondrial nucleoids are left behind, indicating that the cystic bulge does not completely eliminate nucleoids from *tam^RNAi^* spermatids (Figure 2D,E and 3C). To examine how Tam and EndoG might work together to facilitate mitochondrial DNA elimination, we expressed *bam>tam^RNAi^* in the *EndoG* mutant. The flies with both a knockdown of Tam and mutation of *EndoG* exhibited a phenotype that was not obviously different from that resulting from *bam>tam^RNAi^* alone: the mitochondrial nucleoids persisted through late elongation and individualization stages and were only partially collected in the cystic bulge (Figure 3). The failure of the backup mechanism to eliminate the genomes following *tam* RNAi might indicate that this process also requires Tam, however it is also possible that this mechanism, which appears to still function to some degree, is overwhelmed by abundant persisting genomes following Tam knockdown. In any case, the high abundance and prolonged persistence of nucleoids following Tam knockdown, suggests that the DNA polymerase plays a more crucial role in mitochondrial genome elimination than EndoG.

**Figure 3.**
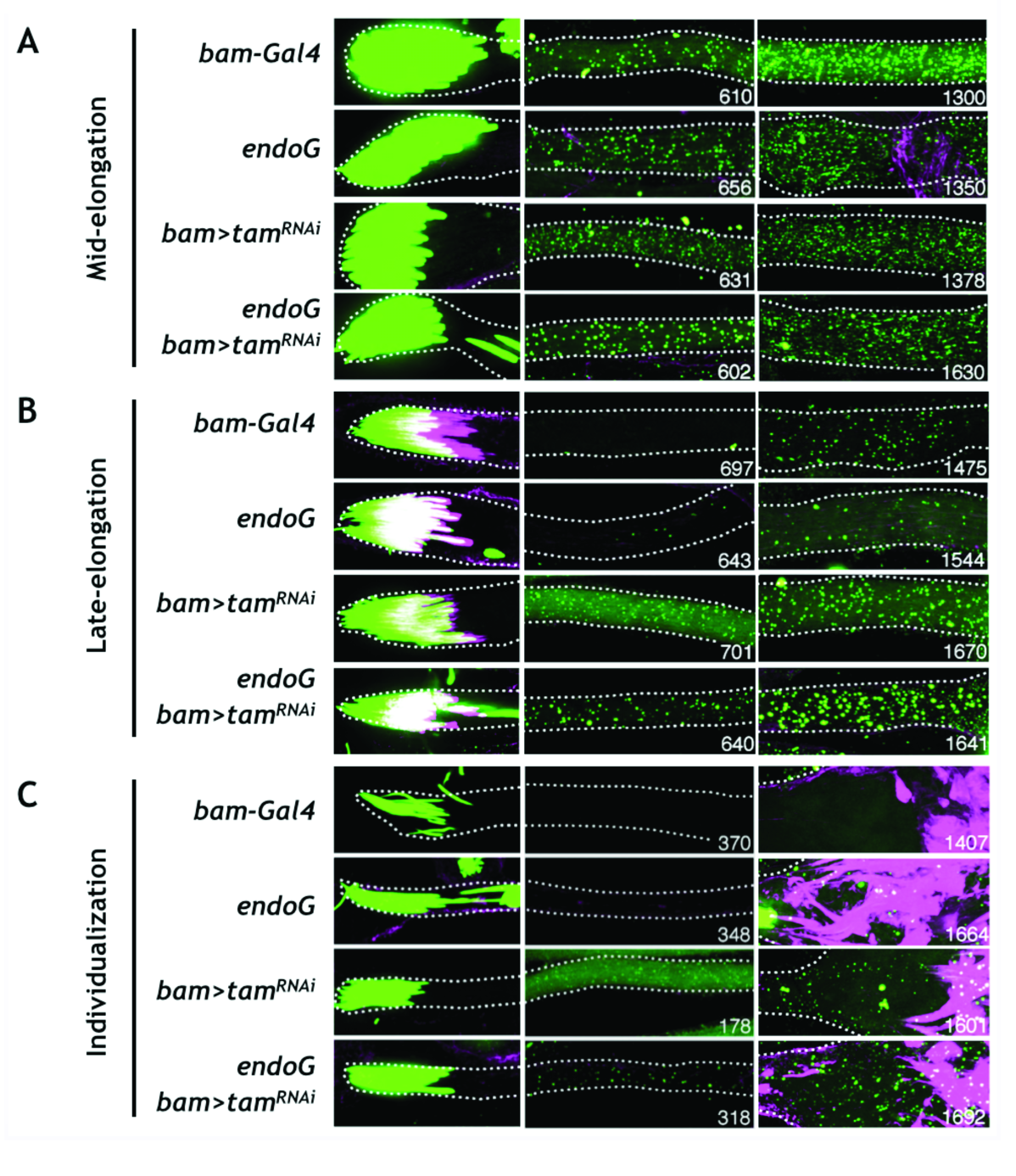
Genetic interactions between *tam* and *EndoG*. **(A-C)** Spermatid bundles of the indicated stage and genotype stained for DNA (PicoGreen: green) and actin (Phalloidin: magenta). Numbers indicate the distance of each image (µm) from basal tip of the spermatid.

### Recruitment of Tam to mitochondrial nucleoids is coordinated with nucleoid destruction

To explore how Tam might contribute to mtDNA elimination, we expressed a C-terminally GFP tagged version of Tam inserted as a transgene that includes the endogenous regulatory sequences and promoter [38], and examined its function (Figure S1) and its localization at different stages of spermatogenesis. As expected for a mitochondrial protein, Tam-GFP was localized to nebenkerns in pre-elongation sperm cysts (Figure S2) and was visualized within the large mitochondria as the nebenkern unfolds and the mitochondria extend alongside the developing axoneme (Figure 4).

At high resolution, Tam-GFP is seen in puncta (Figure 4 and Figure S2). Antibody staining of endogenous Tam showed similar puncta (Figure S3). We assessed co-localization of Tam-GFP puncta with the foci of DAPI staining (nucleoids) visually (e.g. Figure 4A-C). We also quantified co-localization by enumerating pixels having above background staining for GFP or for DAPI or for both in different regions of sperm tail bundles (Methods) (e.g. Figure 4D-F). Both DAPI and GFP staining extended distally as the tails elongated. As expected, the number of DAPI-positive pixels declined in the more elongated (older) sperm bundles as the nucleoids were eliminated. The decline began proximally (Figure 4D).

**Figure 4.**
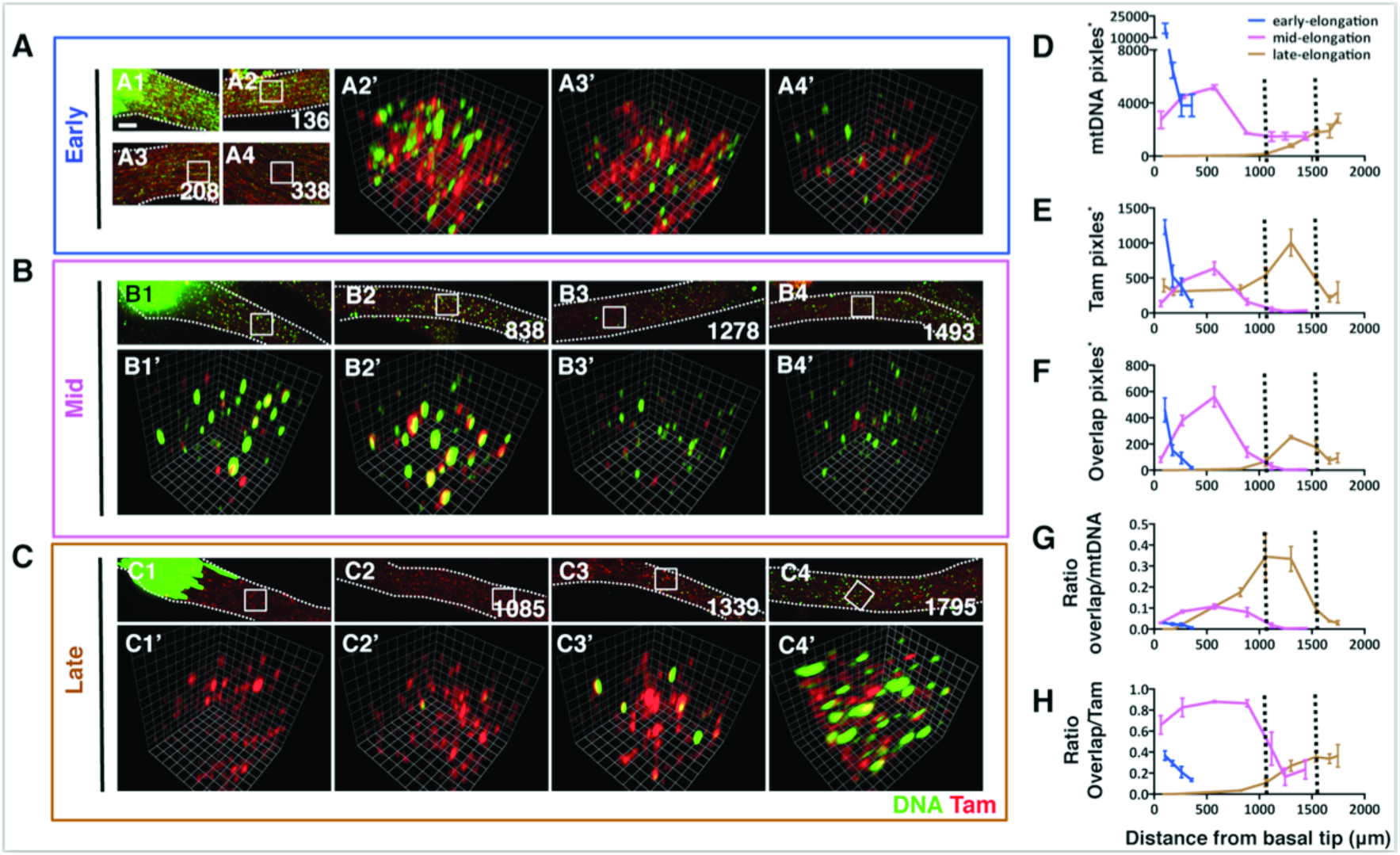
Tam associated with nucleoids prior to their destruction. **(A-C)** Images of elongating spermatid bundles (526,1570 and 1846µm in length, respectively) showing DNA (DAPI: green) and Tam(red, indicates Tam-GFP). Numbers indicate the distance of each image (µm) from basal tip of the spermatid. Areas highlighted in white squares (6x6µm) are shown in magnified 3D opacity views. Scale bars are 5µm. **(D-E)** Pixel counts having above threshold staining for mtDNA and/or Tam within a volume beneath each of three 6x6µm areas such as those marked in A-C were averaged to give the values and SD at the positions indicated. **(F)** The overlap pixel counts represent those pixels with above threshold staining for both GFP and DAPI as calculated by MATLAB program described in Materials and Methods. **(G-H)** Ratio of overlap/mtDNA and overlap/Tam are generated by dividing the number of overlap pixels in F to mtDNA or Tam pixels along the sperm bundles. The early-, mid- and late-elongation stages are indicated in blue, magenta and brown, respectively. * indicates that pixel-number is a sum from imagining planes through the depth of the tail bundle.

We reasoned that if Tam were directly involved with mtDNA elimination, it would be associated with nucleoids at the time of their disappearance, which occurs in a wave moving from the proximal end of the tail to the distal end during late elongation of the sperm tail ([7] and Figure 3). Furthermore, if the recruitment of Tam to the mitochondrial genome is part of the trigger initiating mitochondrial genome degradation, Tam should not be associated with mitochondrial DNA early during sperm tail elongation, before elimination starts. Visible nucleoids and Tam foci were abundant in the short tails of early elongating cysts, but there was very little overlap (Figure 4A). Quantification of pixels staining for DAPI and Tam-GFP showed a high density of staining pixels and a substantial number of dual-staining pixels in the most proximal regions of the tails (Figure 4F). However, dual-staining pixels represented a small proportion of staining pixels (Figure 4G and 4H) and calculation of chance coincidence suggests that a random distribution of the locally high staining would give the observed level of dual-staining pixels (Figure 4D and E and Figure S4). From this we conclude that there is little if any association of Tam foci with nucleoids early during tail elongation.

When the sperm tails extended to about 75% of final length (mid-elongation, 1570um in Figure 4), about the time that nucleoid destruction initiates in the most proximal regions of the tail, we observed frequent overlap of the DAPI and the Tam-GFP foci (Figure 4B). Pixel quantification revealed a high proportion of dual positive pixels peaking at more than 85% of Tam-GFP positive pixels also staining for DAPI (Figure 4H), a level of overlap well above chance (Figure S4). In late elongation bundles, DAPI staining foci are absent proximally—not surprisingly, this change is associated with a decline in dual staining pixels, but it is notable that there is also a decline in total GFP positive pixels suggesting a redistribution of Tam distally in conjunction with the distal progression of the wave of destruction. A high proportion (33%) of DAPI positive pixels co-stain with GFP specifically in the region across which DAPI foci decline (rectangle in Figure 4C3′ and D-G). Toward the most apical end of the tail (1747/1864µm), only 3% of the DAPI positive pixels are GFP positive (Figure 4G), consistent with a lag in the disappearance of the most distal nucleoids. These observations reveal a traveling wave of Tam-GFP localization to nucleoids that parallels the timing of the wave of disappearance of nucleoids. The timing of this association between Tam and nucleoids suggests that requirement for Tam reflects a direct involvement of the protein in the elimination process.

### The exo-domain is dispensable for Tam mediated mtDNA elimination

Like many DNA polymerases, Tam has a 3′-exo-domain that is generally thought to have a proofreading function during replication but which can also degrade DNA from exposed 3′ ends [33]. We expected that this nuclease activity would underlie the contribution of the polymerase to nucleoid elimination, and envisaged a collaboration of endo- and exonucleolytic activities in genome elimination. The exo-domain is highly conserved and D263 is reported to be important for exonuclease activity across species [32,39]. Since homozygous exo-nuclease deficient *tam* mutants are not viable [39], we devised a test based on an assay of the ability of Tam or a mutant form of Tam to rescue the defect in paternal mtDNA elimination caused by *tam^RNAi^*. By altering codon usage, we made a version of *tam* that is resistant to *tam^v106955^RNAi* (Figure 5A). We made a transgenic line expressing a wild-type version of this RNAi-resistant gene, *UASp-tam^wt^-resistant*, and one expressing an exo-nuclease deficient version, *UASp-tam^D263A^-resistant* (Figure 5A). We use RT-PCR with primers specific for the endogenous gene to follow KD of the endogenous transcript. The expression of RNAi resistant *tam* transgenes did not interfere with RNAi knockdown of the endogenous gene (Figure 5B). We then tested the ability of RNAi resistant transgenes to rescue. Expression of the control *UASp-tam^wt-^resistant* transgene completely rescued the defect caused by *tam^RNAi^*. That is, mtDNA was eliminated at the late elongation stage and no mtDNA could be visualized in cystic bulges during individualization (Figure 5C,E). Surprisingly, we observed a similar rescue using *UASp-tam^D263A^-resistant* (Figure 5D and 5E). Thus, the D263A mutation does not compromise the ability of Tam to contribute to the DNA elimination program indicating that the exo-nuclease activity of Tam is not required for nucleoid elimination (Figure 5D and 5E).

**Figure 5.**
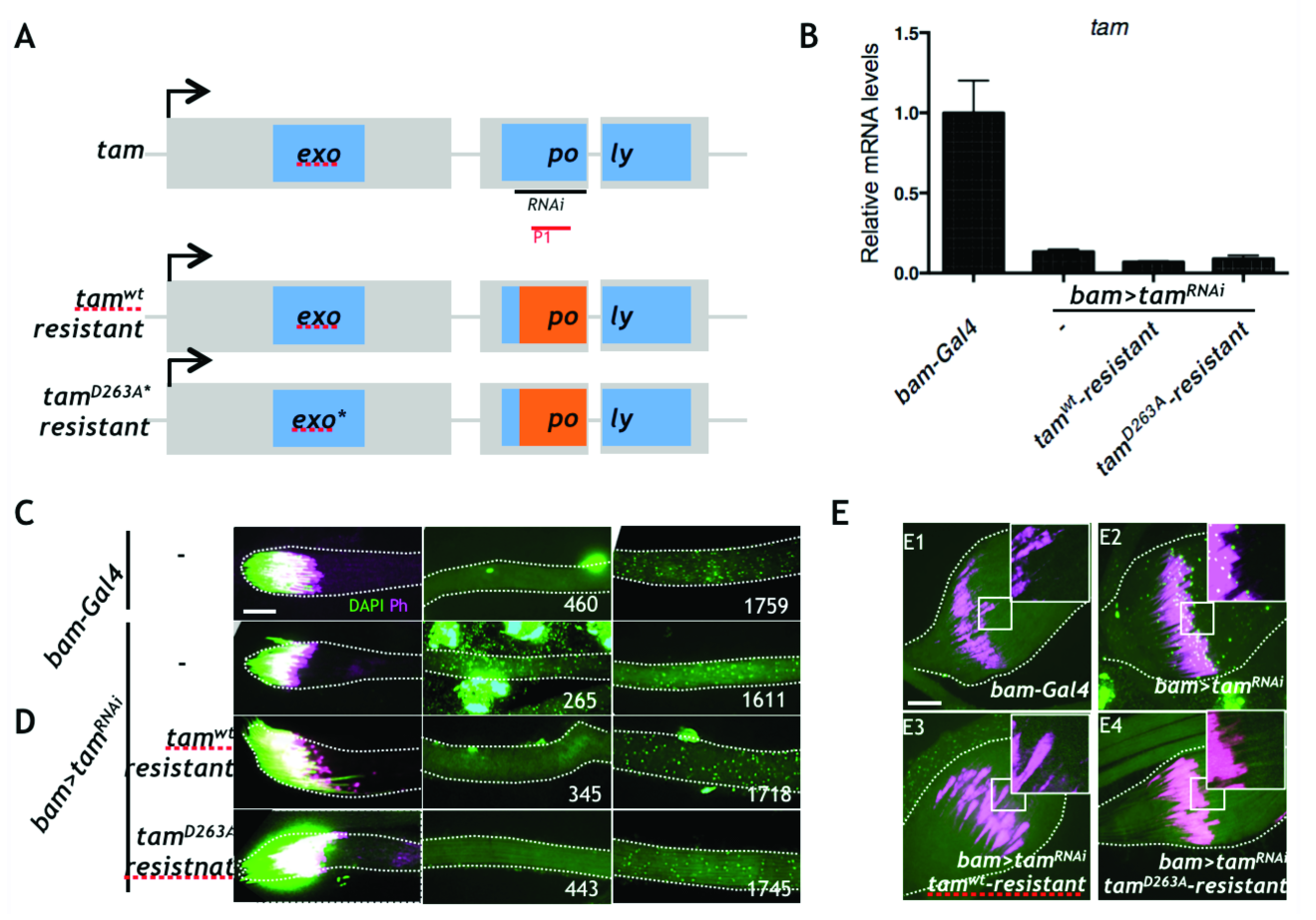
Tam exo-domain is dispensable for mtDNA elimination. **(A)** Schematic of the *tam* gene showing the region targeted by *tam^v106955^RNAi* and *tam* rescue constructs (wild type and exo-domain mutant) containing an RNAi resistant sequence (orange). P1 (red line) indicates the region amplified for RT-qPCR assays of the endogenous *tam* RNA. **(B)** qPCR specifically measuring endogenous *tam* mRNA in adult male testis with different genotypes. Error bars represent the standard deviation for three independent experiments. **(C-D)** Late elongation spermatid bundles stained for DAPI (DNA) and Ph (Phalloidin, actin-cone). Numbers indicate the distance of each image (µm) from basal tip of the spermatid. **(C)** *bam-GAL4* control, 1945µm **(D)** *bam>tam^RNAi^* alone, and with *tam^wt^-resistant*, or with *tam^D263A^-resistant*, (total cyst length 1904, 1986, and 1769µm, respectively). **(E)** Cystic Bulge stained for DAPI (DNA) and Ph (Phalloidin, investment complex). Genotype for each cystic bulge is indicated at the bottom of each picture. The empty boxes (10x10µm) with white border are magnified and showed at the right corner. Scale bars are 10µm.

### Persistent mtDNA in sperm after Tam knockdown

The above findings show that the knockdown of Tam interferes with the normal developmentally programmed disappearance of mtDNA, but we wanted to test whether the persistence of mtDNA extended to mature sperm and whether this persistence would lead to transmission of paternal mtDNA. We collected individualized sperm from the male sperm storage organ, the seminal vesicle. We found no DAPI foci in the tails of control sperm, while sperm from *bam>tam^RNAi^*-expressing flies had obvious DAPI staining foci (Figure 6A).

To examine sperm at an even later stage and to quantify the level of remaining mtDNA, we mated males having wild type mtDNA (*mt:ND2*) to females having a 9 base pair mtDNA deletion, *mt:ND2^del1^*, that is not amplified by wild type specific PCR primers, dissected out the female sperm storage organ from these mated females and used qPCR to quantify the amount of mtDNA transferred from the male [7]. Consistent with previous observations, we did not detect male mtDNA in storage organs from females mated to control males (*bam-GAL4*, Figure 2C). However, we detected male mtDNA in storage organs from females mated to *tam-RNAi* expressing males (Figure 6B).

**Figure 6.**
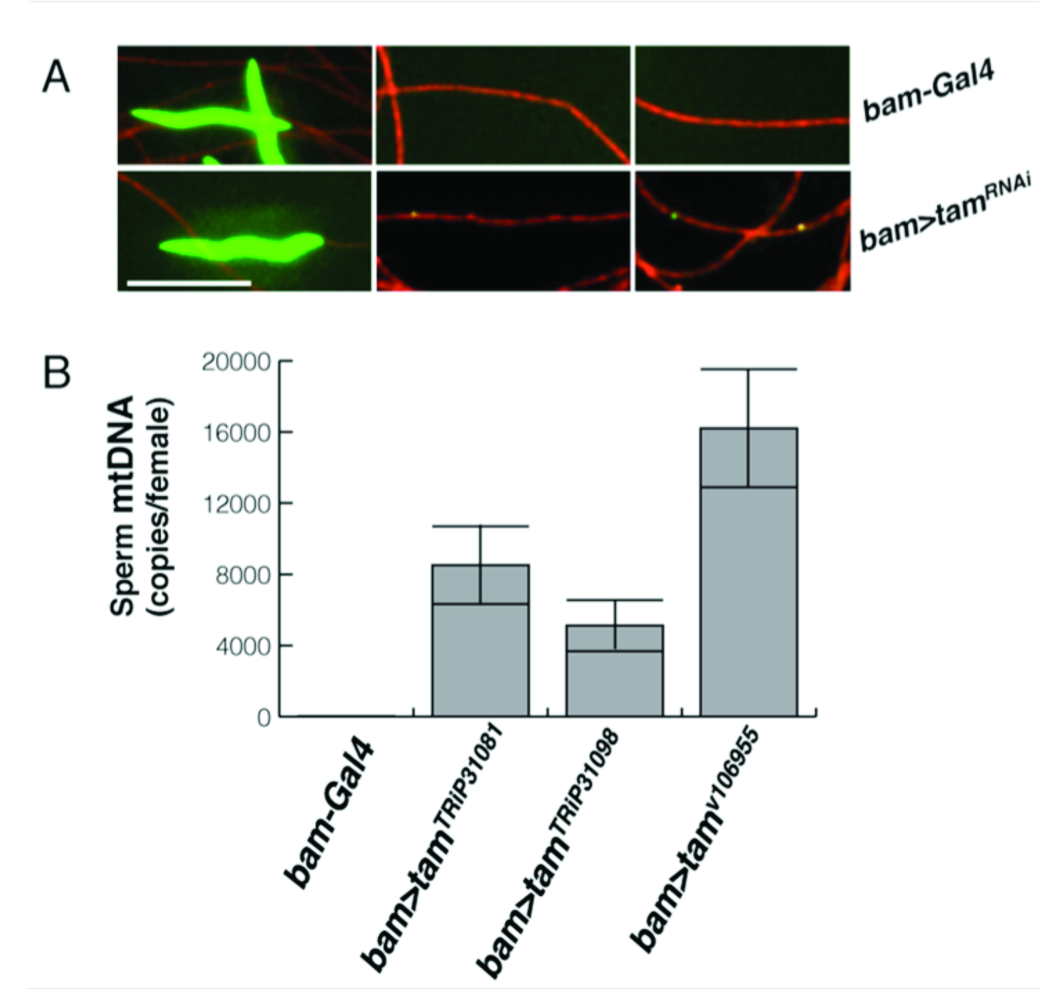
*tam* knock-down prevents mtDNA elimination from the sperm. (**A-B**) Sperm collected from seminal vesicles. DAPI (green) shows DNA, and mtSSB-RFP (red, see Materials and Methods part) marks the mitochondria of the sperm tail. Because the sperm are individualized, if there were no loss of mtDNA the density of nucleoids a tail is expected to be 1/64^th^ of that seen in the sperm tail bundles of developing cysts. The length of the tails and tangling of the sperm made tracing of individual sperm impractical, so heads (left) and tails (middle and right) are independent. Scale bar is 6µm. **(A)** *bam-GAL4*. **(B)** *bam*>*tam*^*RNAi*^. **(C) Quantification of male mitochondrial genomes** (*mt:ND2*) transferred to the sperm-storage organs of *mt:ND2^del1^* females following matings with control males (*bam-Gal4*) or males expressing one of three *tam*^*RNAi*^ constructs. qPCR with *mt:ND2* specific primers measured the amount of male-derived (sperm) mtDNA in mated female sperm-storage-organs, which can store up to 1000 sperm. The number of sperm mtDNA per female storage organ is plotted.

The presence of mtDNA in the sperm contained in the female’s sperm storage organ suggests that the persisting DNA will be transferred to fertilized eggs along with the mitochondria. However, a qPCR measure of the average amount of paternal mtDNA in a collection of 0-3 hour old embryos fertilized by sperm from Tam knockdown flies was about 0.3 copies per early embryo (Figure S5). This measured level is above the control level (0.03 genomes per embryo), but it is much less than expected if each sperm where to deliver its load of multiple genomes. However, the genomes initially delivered might have been rapidly eliminated since the paternal mitochondria themselves are eliminated during early embryonic development [11,12]. Apparently, the embryonic elimination of paternal mtDNA is rapid, or the sperm are heterogeneous in their mtDNA content and only sperm with little or no remaining DNA are responsible for the majority of the fertilization.

## DISCUSSION

We report an unexpected role of the mitochondrial DNA polymerase in eliminating mitochondrial genomes during spermatogenesis. In addition to influencing our view of the functions of a DNA polymerase, the finding reinforces previous indications that the nuclear genome actively restricts the transmission of the mitochondrial genome [7]. It has long been recognized that the differences in the routes by which nuclear and mitochondrial genes achieve successful transmission create evolutionary conflicts, and that cooperation is more like a standoff between combatants [18]. An unrelenting battle with participants constantly struggling to outdo each other accelerates evolution in a “Red Queen” process in which a change in one genome selects for an adjustment in the other [40,41]. But when each genome also relies on the other, this process can also be viewed as a search for stability, since only more stable standoffs will persist. We propose that use of POLG (Pol gamma) in mitochondrial genome elimination contributes to stability because mtDNA cannot easily avoid destruction without also avoiding replication.

There is no direct information from other organisms to indicate whether POLG is used widely for DNA elimination programs that enforce uniparental inheritance. However, infectious mitochondria-localized plasmids found in some plant and fungal species can escape paternal mitochondrial genome elimination and are able to spread through biparental inheritance [19,20]. Many of these plasmids encode their own DNA polymerase, and in some plasmids, the polymerase is the only encoded gene [19]. By encoding their own DNA polymerase, these plasmids would not require POLG to replicate and could avoid interacting with POLG altogether, allowing them to escape a POLG-dependent destruction mechanism.

We have considered several ways in which POLG-dependent destruction could occur. We started with the simple idea that the 3′-exonuclease activity of Tam would be directly involved in destroying the nucleoids. However, we demonstrated that a mutant form of Tam lacking 3′-exo function still promoted nucleoid elimination. Alternatively, Tam may sensitize the mtDNA for destruction by another nuclease, or recruit another nuclease to the mtDNA. In either case, mtDNA and a destructive nuclease could stably co-exist within mitochondria until mtDNA associates with Tam. During *Arabidopsis* pollen development, a nuclease degrades mtDNA in generative cells to prevent paternal inheritance [42]. A homologous nuclease is targeted to mitochondria during cucumber pollen development, but mtDNA in cucumber generative cells remains unharmed and is paternally transmitted [43]. Apparently, the presence of a nuclease is not sufficient for elimination. An attractive possibility is that cucumber generative cells down regulate POLG-like activity while *Arabidopsis* generative cells posses it. Note here, that in this scenario, escape of the elimination program is due a change in the nuclear program, and is not under the control of the mitochondrial genome.

Despite remaining mysteries, our findings uncovered a new action of DNA polymerase — the elimination of mitochondrial genomes. By controlling the different outcomes — replication or destruction— the nuclear genome acquires a special “power” over mtDNA. This power may be needed to counter a selfish drive that promotes ever more potent mtDNA replicators [44]. It is inevitable that different mitochondrial genomes in an organism compete with each other for transmission. In this context, genomes that evade limits and replicate more effectively than their sibling genomes will thrive. The evolution of juggernaut replicators could be to the disadvantage of the host and result in another genetic conflict between nuclear and mitochondrial genomes. Genome elimination is a tool that the nucleus could use to contain the unruly behavior of mitochondrial genomes.

DNA is viewed as a stable repository of genetic information, but labeling experiments in rats revealed a steady state of mtDNA replication and destruction in non-growing postmitotic tissues [45]. If turnover is similar in humans, the mitochondrial genomes of our quiescent tissues could be replaced 10,000 times over a lifetime. Surprisingly, the basis for this turnover is not known. Its likely importance is highlighted by a group of mitochondrial based metabolic diseases. Mutations in nuclear genes regulating mtDNA replication and deoxynucleotide pools result in disease phenotypes marked by accumulation of mtDNA defects, particularly deletions and reduced copy number [25,26]. If these mutations reduce replication, they should show reduced mtDNA copy number expansion during extensive embryonic growth. Instead, the defects are predominantly postnatal, and often later in life. Some of the mutations, particularly dominant mutations in the mitochondrial DNA polymerase that give rise to progressive external ophthalmoplagia (POE) [27], might be candidates for alleles that enhance DNA elimination rather than alleles compromising replication. Mitochondrial DNA deletions and depletion that accompany aging suggest imperfections in the dynamic maintenance of mitochondrial DNA in “quiescent” cells. It will be interesting to see whether nuclear-programmed elimination of mitochondrial genomes like that operating during *Drosophila* spermatogenesis has a more widespread impact on our biology.

## MATERIALS AND METHODS

### Antibody and Fluorescence microscopy

For immunization, DNA fragments coding for *Drosophila* Tam amino acids 30 - 653 was cloned into pET28a plasmid using NotI and NdeI sites. Recombinant plasmid was transformed into the BL21(DE3) competent *E.coli* cells and His-Tag fused protein was purified using the Ni Sepharose (Sigma-Aldrich) manufacturing instruction. The subsequent rabbit immunization with the His-Tagged Tam was carried out by Pacific Immunology. For antibodies purification, the same DNA fragment was cloned into pET41a plasmid using SpeI and NotI sites. The resulting GST-fusion protein (Tam-GST) was expressed in bacteria, trapped on a Sepharose-GSH resin, and crosslinked to this resin using disuccinimidyl suberate (Thermo Fisher). The antibody was purified by binding to this Tam-GST column followed by elution at low pH. Eluted antibody was neutralized and diluted for staining.

DAPI or PicoGreen/Phalloidin and antibody staining and imaging of fixed spermatid cysts and individualized sperm from seminal vesicle were performed as previously described [7]. Images were acquired using a spinning disc confocal microscope, processed with Volocity 6 Software. The lengths of spermatid cysts were measured using Volocity 6 Measurements module. To illustrate the recruitment of Tam to mitochondrial nucleoids during sperm development (Figure 4), the Volocity 3D opacity module was used to generate a perspective image from a collapsed z-stack.

### Quantitative image analysis to measure expression level and spatial overlapping

To calculate the number of Tam-GFP, DAPI or dual staining pixels, at a given position along each analyzed spermatid bundle three nearby 6 × 6μm squares were chosen for each data point. Within each area, a full stack of images representing the depth of the sperm tail bundle was analyzed using MATLAB. The program identified and enumerated pixels having a signal above threshold in either or both DAPI and GFP channels. The numbers of such pixels for each of the three areas analyzed at each position were averaged and plotted versus position along the bundles as shown in Figure 4D-4H. The source code will be provided on request.

To calculate the theoretical expectation for coincidental spatial overlap (Figure S4A), the following method was used: assuming independent and random distribution of mtDNA and Tam-GFP pixels, the theoretical expectation of spatial overlap fraction of these two channels is governed by (v_DAPI_/v)* (v_GFP_/v), where v_DAPI_, v_GFP_ are pixels measured in each stack of image with mtDNA or Tam-GFP signal respectively (Figure 4D and Figure 4E) and v denotes the total number of pixels in the same stack of image. In contrast, the actual spatial overlap fraction (Figure S4B and Figure 4F) is given by the ratio v_overlap_/v, where v_overlap_ is the measured number of pixels showing dual staining.

### qPCR measurement of mtDNA copy number in sperm and paternal mtDNA copy number in embryos

*mt:ND^del1^* females were mated to *mt:ND2*, *bam-GAL4* (control) or *mt:ND2*, *bam-GAL4/UAS-tam^RNAi^* males. *mt:ND2* (male-derived) copy number in *mt:ND2* female sperm storage organs or 0-3 hour old embryos was measured as previously described [7].

### Measurement of Tam mRNA in whole testis

For each experiment, 20 testes were dissected and pooled in PBS. Total RNA from testes of the stated genotype was isolated using Trizol reagent (Invitrogen) and further cleaned with TURBO DNase (Ambion). cDNA was prepared using the SensiFast cDNA Synthesis Kit (BIOLINE). Quantitative PCR (qPCR) was performed on a real-time detector machine (STRATAGENE-MX300p) and results were analyzed and graphed with the software GraphPad Prism6. All qPCR values are the mean of three independent experiments after normalization. Gene-specific primers used for qPCR are listed below (5’-3’): tam-forward, 5’-ATTGACCAGCTCCGTAGATC; tam-reverse, GGGTTATAGACTCGTTGGTG; Rp49-forward, ACGTTGTGCACCAGGAACTT; Rp49-reverse, TACAGGCCCAGATCGTGAA.

## AUTHOR CONTRIBUTIONS

SZD made the initial discovery, SZD and ZY planned and performed experiments, ZY, SZD and PHO’F contributed to conception, interpretation and the writing of the manuscript, and PHO’F supervised and obtained funding.

## ACKNOWLEDGEMENTS

This work was supported by the NIH ES020725 and GM120005 funding to P.H.O’F. S.Z.D. was supported by a Larry L. Hillblom Foundation award. We thank Juan Guan and Bo Huang for kindly providing the MATLAB source codes and helping with image analysis. We gratefully acknowledge the Bloomington Stock Center, the Vienna Stock Center, Minx Fuller, and Hong Xu for stocks. We thank members of the O’Farrell laboratory for discussion.

